# Modelling *in vitro* mycelial growth, sporulation and spore germination of *Pseudocercospora musae* under different illuminance levels

**DOI:** 10.1101/795450

**Authors:** Djalma M. Santana-Filho, Milene C. da Silva, Zilton J. M. Cordeiro, Hermínio S. Rocha, Francisco F. Laranjeira

## Abstract

Banana is one of the most produced fruits in the world. Many diseases infect the culture and yellow sigatoka is one of the most important. Light may interfere in the pre-penetration parameters of the fungus. The aim of this study was to evaluate the action of light on aspects of the life cycle of the causal agent. The action of light was tested in *vitro* on the mycelial growth, sporulation and germination of the fungus. For the mycelial growth 10 colonies were transferred to each Petri plate, by evaluating the green weight of a plate at each moment, in mg, at zero, 15, 21, 28, 35 and 42 days of cultivation according to different illuminance levels. Under the same conditions, the sporulation was also quantified at 5, 7, 11, 13 and 14 days. For germination, equal volumes of a spore suspension were placed in small glass containers, which then were set in mini-shades (different illuminances). Every 1h, lacto-phenol was added to a glass from each environment paralyzing the growth, at the same time, illuminance measurements were made with a light meter. The data obtained from sporulation (linear relation) and germination (positive exponential behavior) were significant for the illuminance levels tested.

**Significance and Impact of the Study:** Yellow Sigatoka disease, caused by *Pseudocercospora musae*, is still one of the most importante banana diseases in Brazil. Its control demands the use of fungicides. New environmental-friendly control methods should be developed. Therefore, the behaviour of both fungus and plant-fungus interaction should be known. We modelled the in vitro behaviour of *P. musae* in function of light intensity. Our results can help to develop shading strategies to control yellow Sigatoka.

## INTRODUCTION

Brazil is one of the largest banana producers in the world (FAO, 2012). However, despite the prominent position in world production, the country still faces serious problems of phytosanitary management (Cordeiro and Kimati, 2005).

Yellow Sigatoka, caused by *Pseudocercospora musae* (previously known as *Mycosphaerella musicola*) is one of the main diseases of the banana crop in the regions where the environmental conditions favor it, and it is found in all banana-producing regions, with some exceptions (Cordeiro and Matos, 2001, 2003), adapting well to cooler areas, with altitudes above 1200 meters (Jácome, 2002). The damage from both Yellow and Black Sigatoka (*Pseudocercospora fijiensis*, previously known as *Mycosphaerella fijiensis*) diseases can be higher in plantations without proper control, reaching 100% if the microclimate is favorable, and the fruits produced are of no commercial value (Cordeiro, 2004).

Based on origin, semi-shaded environments are the most appropriate for banana cropping (Favreto *et al*, 2007). The low incidence of Black Sigatoka disease and low severity in places where the brightness is reduced support this hypothesis (Beltrán-Gracía *et al*, 2014; Norgrove, 1998; Norgrove and Hauser, 2012). In environments where there is only 30% light incidence there is a significant delay in yellow sigatoka infestation, with a low number of necrotic leaves, and the disease development time is four days longer in about 70% of light environments. Despite this, the first lesions appear in equal times in both conditions (Dold *et al*, 2008).

The light effects can be direct or indirect intensity, with further damage occurring in treatments with higher luminosity levels in Black Sigatoka disease (Norgrove *et al*, 2012). In high-density planting systems, the incidence is not affected, but the severity of the disease is reduced by 18% (Emebiri and Obiefuna, 1992).

Agroforestry systems have significantly reduced the severity of yellow and black sigatoka, possibly due to reductions in UV radiation that is necessary for sporulation and the release of the fungus spores (Schrotz *et al*, 2000).

Significant results show a greater number of spores produced in continuous light than in 12 hours of light or total darkness. In addition, light pulses of five minutes for four days can induce sporulation of the pathogen. This demonstrates that the spectrum of light has a significant effect on the formation of *P. fijiensis* conidia (Sepulveda *et al*, 2009). Albuquerque (1993) found similar, but not significant, results between 12 hours of total light and total darkness.

For Abreu (2000), sporulation begins after three days of cultivation, and may fluctuate over time for *P. musae*. The number of *P. fijiensis* colonies is smaller in the dark when the plates are sealed for 21 or 14 days (Etebu *et al*, 2005).

Some studies on the interactions between the banana – *P. fijiensis* show that toxin production is associated with the presence of light (Lepoivre et al. 2002), others that ultraviolet radiation (UV) is a limiting factor in the production of *P. musae* ascospores, justifying the low sporulation (Jones, 2002).

According to some authors, shading reduces cercosporin activity, involved in the pathogenesis, which is dependent on photosensitization (Daub and Ehrenshalf, 2000; apud Gasparoto *et al* 2010; Daub *et al*, 2005). However, studies report that the pigments contained in the mycelium and secreted in the culture medium of *P. fijiensis* are melanins that absorb visible light and act as photosensitizers, which may generate O_2_, suggesting that this should be studied further as a potentially important contributor to the progression of black Sigatoka in bananas and plantains (Beltrán-Gracía *et al*, 2014).

Chemical and morphological differences were determined between three species of fungi in the *Pseudocercospora* complex. Preliminary data showed a chemical toxin production route, with minor differences between fungi (Stierle *et al*, 1991).

Plants under the shading system present low disease severity. This may be explained by reduction in dew or light incidence (Cordeiro, Matos and Kimati, 2005). The shadow produced by the dominant extracts is an important reducer of the damage caused by yellow sigatoka (Vivan, 2002).

The aim of this study was to investigate the behavior of the pre-penetration parameters of the fungus under different illuminance levels. This independent variable is the limit of the ratio of the luminous flow received by the surface around a point considered for the surface area when it tends to zero. It is not distributed uniformly in all points of a given area, so the average is often adopted to determine whether the values are suitable for the performance of certain activities (Bormann, 2003). The hypothesis tested was that higher illuminance intensities induce greater mycelial growth, sporulation and germination of the pathogen’s conidia.

## MATERIALS AND METHODS

The experiments were conducted in the Plant Pathology Laboratory at Embrapa Cassava and Fruits (Cruz das Almas, Bahia). The growth parameters of colonies, sporulation and germination of the fungus *Pseudocercospora musae* were evaluated under different illuminances.

### Isolate collection and sporulation induction

The fungus was isolated from the leaves of the banana plant ‘Pacovan’ that present characteristic symptoms of yellow Sigatoka. The collections were made in June 2010, in the town of Laranjeiras (Muritiba, Bahia). Both the collection and the sporulation of the isolate followed the methodology described by Cordeiro, Rocha and Araújo (2011). Symptomatic leaves were collected and placed in a moisture chamber until sporodochia emergence. These structures were observed under a microscope and taken to agar media. The colonies observed after a few days were picked and transferred to malt medium and after growth were macerated and re-passed to sporulated in V8 media.

### *In vitro* mycelial growth

In this experiment the *in vitro* mycelial growth was evaluated in function of the illuminance measured with a light meter. Thirty-six plates were prepared with V8 culture medium and 10 *P. musae* colonies were placed per plate, selected with the highest uniformity possible. The treatments were four levels of illuminance 5383 Lux; 110 Lux; 10.2 Lux; 2 Lux. At time zero (control) a total of 80 colonies, 2 plates per treatment, were weighed on precision scales to obtain the green weight. The average green weight of all the treatments resulted in the data for that time. The other assessments were made at 15, 21, 28, 36 and 42 days of fungus cultivation when one plate of each treatment was removed from each treatment, and shortly afterwards, the 10 colonies were weighed on precision scales. The weight of the colonies was evaluated at four illuminance levels, at a temperature of 25°C and a 12-hour luminosity photoperiod.

### *In vitro* Sporulation

The experiment consisted of 4 treatments with 3 replications each, and each plate constituted a replication. The treatments were made with different illuminances (3380, 250, 30 and 1 lux) focusing on the 12-hour photoperiod on the plates. Light levels were obtained by placing the plates under boxes manufactured with different types of shading mesh. The evaluations were made at 5, 7, 11, 13 and 14 days after the suspension was sown. Spores were released following the methodology proposed by Cordeiro, Rocha and Araújo (2011). Then, using a Pasteur pipette, two blades were prepared in Newbauer chambers for each treatment, which were taken to the stereoscopic optical microscope to count the number of spores per ml. Two fields of the same were always counted, making a total of four fields observed.

### Spore germination

In this experiment, a rate of 5ml of homogenized suspension, obtained by the methodology by Cordeiro, Rocha and Araújo (2011), were placed in jars for penicillin substances. The bottles were placed in miniature screen versions produced with the following screens types: clarity (58.976 lx), shadow 25% (44.373 lx), shadow 50% (27.328 lx), shadow 75% (24.673 lx), and a structure covered with 3 layers of shadow 75% (4.181 lx). Six bottles per structure were placed in these replicas, with a total of 30 glass jars containing the spore suspension. Every hour, 1 bottle of each treatment was removed and taken to the lab to paralyze the fungus germination with lacto-phenol. At the same time when the bottles were removed, the illuminance in the bottles was verified under the miniatures, 20 measurements were taken with the light meter obtaining the above measurements presented besides the treatments enclosed in parentheses. The glasses were placed in the refrigerator until they were put under a microscope to assess the number of spores germinated in function of the illuminance levels tested. For the counting, 4 blades were prepared, which were placed under the stereoscopic microscope to count the germinated spores. The percentage of germinated spores was obtained from the count of 100 spores.

### Statistics

The data was analyzed by simple linear and exponential regressions. The software BioStat 5.0 was used in all three cases: in which colony growth, sporulation and fungus germination, were the dependent variable and illuminance was the independent variable. The dependent variable for each case was, respectively: weight of the colonies in mg, the number of spores.ml^-1^ and the percentage of spore germination. For the variable sporulation the data passed by a logarithmic transformation of data, because the data was multiplicative.

## RESULTS

### Colony growth

The linear regression model was significantly adjusted to data obtained under high illuminance in incubation conditions (Table 1). However, for low values (as 10.2 and 2 lx) no model has been satisfactorily adjusted. Comparing the regression in both cases, there was no significant difference showing that the linear model is the best to describe the data for mycelial growth. However, the equation for 5383 lx describes 86% of the data whereas the equation for 110 lx describes about 70% of the data.

**Table 1:** Results of the regression analysis for *in vitro* mycelial growth, sporulation and spore germination of *Pseudocercospora musae* at different levels of illuminance.

|  | Equation | r<sup>2</sup> | Regression model |
| --- | --- | --- | --- |
| **Mycelial growth** | $Y' = 3.9897 + 0.0531X$ | 0.696 | Linear |
| **Sporulation** | $Y' = 0.6320 + 0.1343X$ | 0.968 | Linear |
| **Germination** | $Y = 4.014 + 0.0848 \cdot \exp.(-\text{lux}/-13688,6)$ | 0.971 | Exponential |

### Sporulation

The illuminance levels interfered significantly in the pathogen sporulation (p=0.0127). Thus, sporulation increases when the light level increases (Figure 01). 97% of the data analyzed can be explained and adjusted by the linear regression model. The best sporulation averages were obtained under 3380 lx illuminance, which makes this treatment significantly different from the others. The treatment at 250 lx was significantly different from the treatment at 1lx, but there was no significant difference between the treatments at 30 lx and 250 lx.

**Figure 1.**
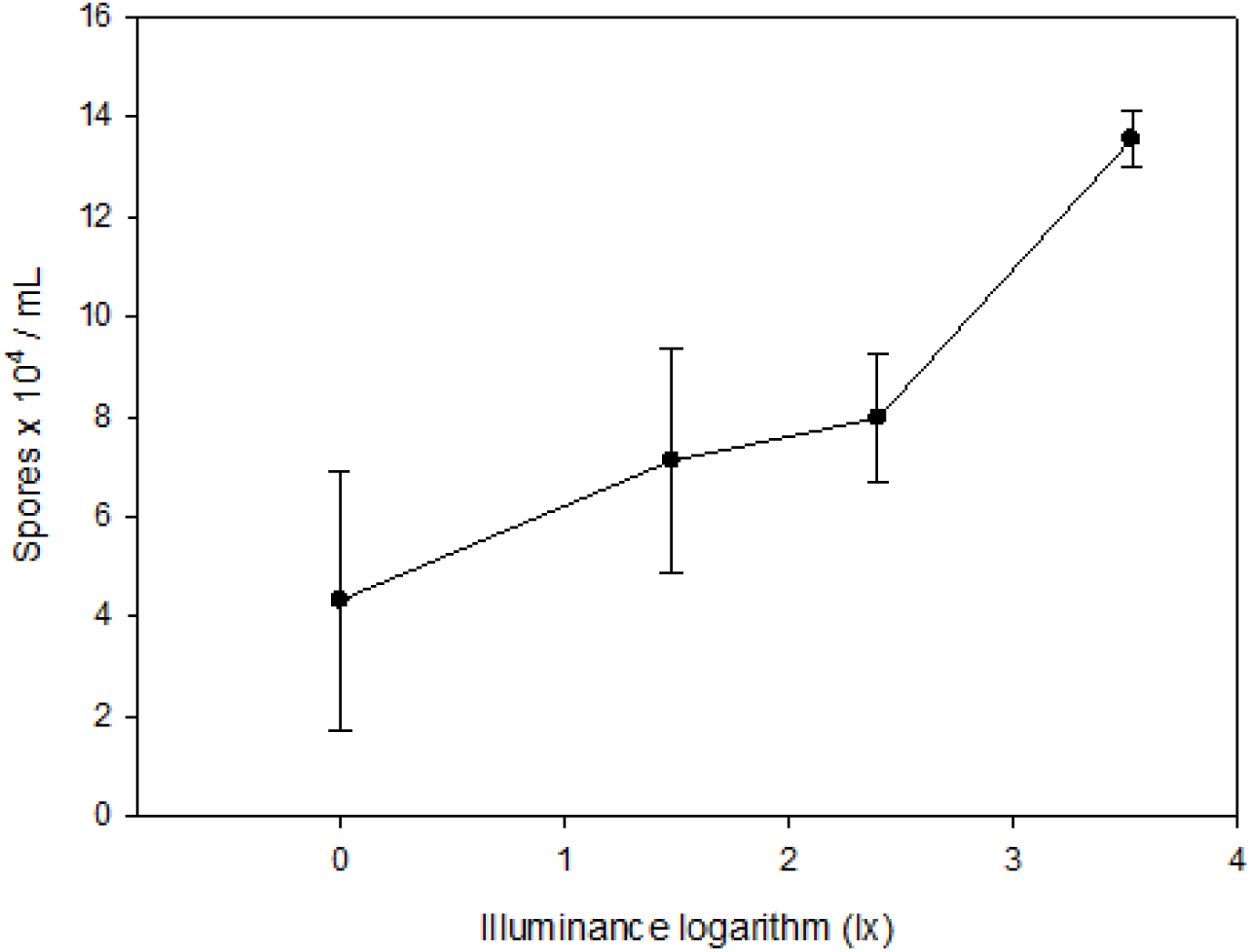
Number of spores per milliliter of spore suspension prepared with the fungus *Mycosphaerella musicola*, cultured on petri plates under illuminances at 3380 lx, 250 lx, 30 lx and 1lx. The data had logarithmic transformation because it is multiplicative data, thus obtaining Log 3380 lx = 3.53, Log 250 lx = 2.40, Log 30 lx = 1.48, Log 1lx = 0.00. The equation was obtained by a linear regression model using the BioStat 5.0 program.

### Germination

The nonlinear model best fitted the germination data and was significant at the level of 5%. There is an exponential relationship between the variables, and the positive effect of the variable on the germination can be verified by its equation. The highest germination percentage was obtained when the spore suspension was under conditions in which the average luminance was 58.975,53 lx (Figure 02). It can also be observed that the germination percentage in 6h is around 10%, demonstrating that the fungus germination of the fungus can take longer than that tested in this experiment.

**Figure 2.**
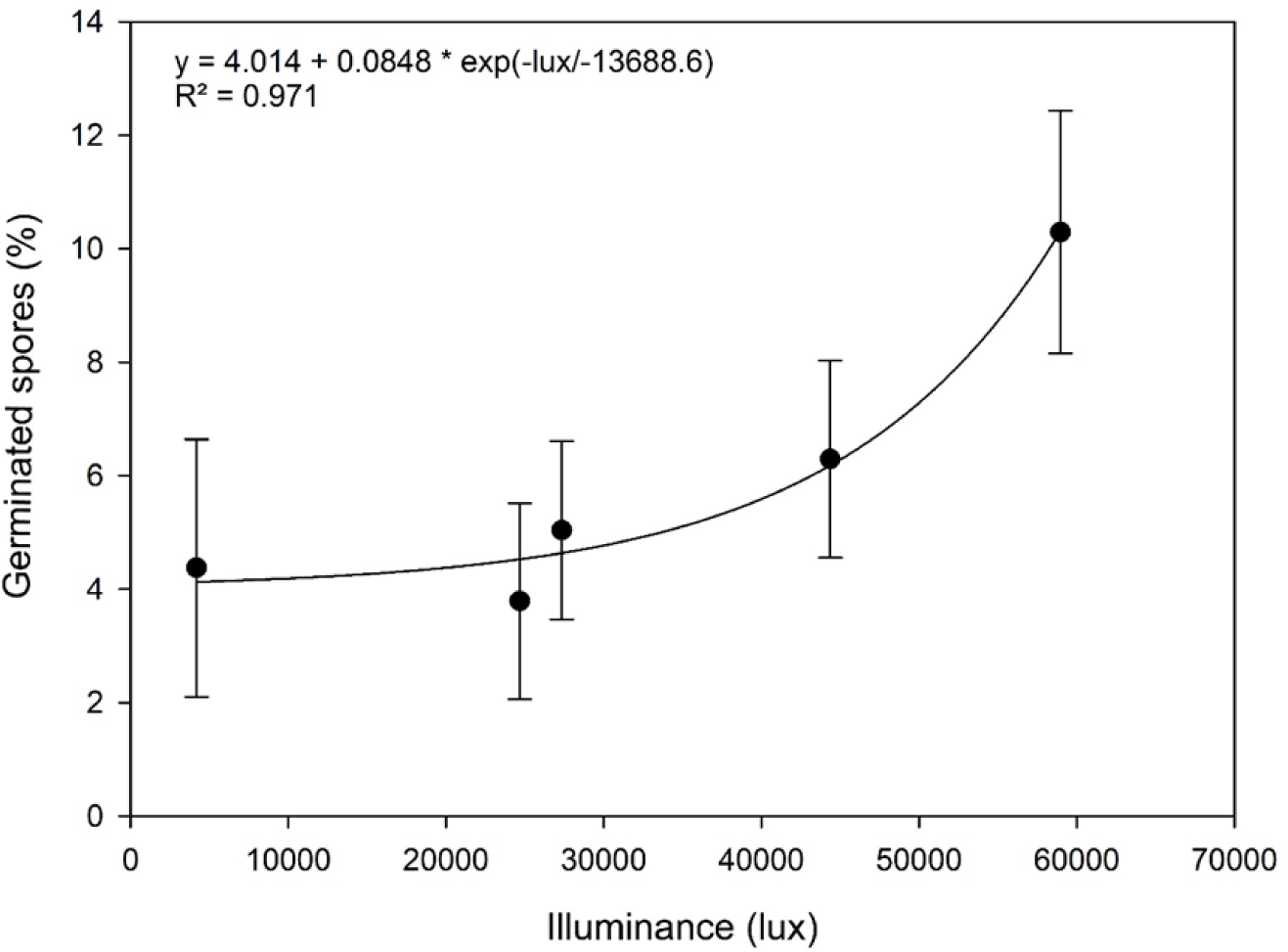
Percentage of *Pseudocercospora musae* spores germinated in vitro under levels of illuminance: 4181.20 lx; 24672.67 lx; 27327.49 lx; 44373.49 lx and 58975.52 lx. The chart presents the model’s equation obtained by an exponential function of three parameters, adjusted by nonlinear regression using the TableCurve 2D program v5.01.

## DISCUSSION

### Colony growth

The results obtained showed no evidence that the mass growth of *P. musae* colonies is affected by illuminance levels, contrary to those obtained by Montarroyos *et al* (2007), who, when evaluating the radial growth of *P. musae* colonies obtained better results in cultures kept in a continuous darkness regime and observed lower averages under the continuous bright clarity regime. Rosa and Menezes (2001) also evaluated the mycelial growth, diameter in millimeters, of different isolates of the fungus in different culture media and pH for 7 days of incubation with 12h light and 12h darkness. Different medias under pH 4.5 induced mycelial growth, with significant differences also for different isolates. However, the evaluation of *P. musae* colony diameter is not the best variable to be quantified in that type of study. The yellow Sigatoka causal agent features vertical mycelium growth and, therefore, the assessment of its mass is more suitable. It can inferred that there is no interference from light on the mycelial growth of the fungus. More detailed studies on the physiology of the fungus in function of light may provide new guidelines related to the action of this factor on the variable mycelial growth and provide plausible interpretations about the action of the fungus in the plant.

### Sporulation

Unlike that observed for colony growth, illuminance had significant effect on *M. musicola* sporulation. The highest sporulation occurs when a greater amount of light reaches the fungus. The highest averages were obtained when the illuminance was higher, around the 10^th^ day after sporulation induction.

The data obtained corroborate those found for the fungi *P. fijiensis* and *P. musae* on a main point. The causal agent of the disease sporulates better in brighter environments (Hanada, Gasparotto and Pereira, 2002), and practically does not produce spores under continuous darkness (Etebu *et al*, 2005; Albuquerque, 1993; Jones, 2002; Lepoivre *et al*, 2002). This was also observed in the study by Sepúlveda *et al* (2009), where there was a higher spore production in the treatment with continuous light, followed by that of the 12-hour photoperiod and then, continuous darkness. Conidia production was very low or nonexistent in dark conditions.

The results concerning the fungus sporulation make it possible to infer that illuminance interferes in the fungus reproduction. Survival may also be affected, if in the absence of susceptible tissue, spores of the fungus and its reproductive structures remain for a few days on decomposing materials. Therefore, under low luminosity, with reduced sporulation, the absence of inoculum can interfere in this parameter of the life cycle of the fungus. With this, the fungus dissemination is also reduced, thus decreasing the likelihood of infections in plants. The secondary cycle of the disease is damaged. The results show that the number of spores produced is less on plates with lower levels of illuminance.

### Germination

For variable germination, the statistical analysis showed that the most appropriate model for germination in function of illuminance was the exponential model of three parameters. Illuminance interferes with the fungus germination affecting the infection, but the low germination percentage demonstrates the need to adjust the methodology applied with regard to its evaluation time. Currently unpublished data on experiments with plants under shade take the hypothesis that the shading reduces the number of lesions on the leaves. This may be related to the low sporulation standards, which decrease self-infections, and the smaller number of stomata presented by acclimated plants under shade conditions (Santana-Filho *et al*, 2012). Some authors studied conidia and ascospors germinations and tub germ growth under different temperatures and moisture wetness (humidity) with *P. fijjiensis* (Jacome *et al*., 2002), but have not studied the effect or interference of light on it or *P. musae*. Our results provide a model about colony growth, sporulation and conidia germination of *P. musae*, the causative fungi of Yellow Sigatoka disease. More information about the physiology of this fungus or similar fungi is required to clarify why and where light interferes in these pre-penetration parameters.

## References

Abreu KCLM (2000) Variabilidade morfológica e patogênica de isolados de Mycosphaerella musicola Leach. M.Sc. Dissertation, Escola de Agronomia da Universidade Federal da Bahia, Brazil.

Albuquerque PSB (1993) Mycosphaerella musicola: Produção de conídios in vitro; Sensibilidade a fungicidas e avaliação da resistência em mudas de cultivares de bananeira (Musa spp. M.Sc. Dissertation, Universidade de São Paulo, Brazil.

Bormann OR (2003) Iluminação natural em salas de aulas e escritórios com uso de prateleiras de luz. M.Sc. Dissertation, Centro Federal de Educação Tecnológica do Paraná, Brazil.

Cordeiro ZJM and Matos AP (2001) Sigatoka-amarela no Norte de Minas Gerais. Simpósio Norte Mineiro sobre a cultura da banana. Porteirinha, MG, p. 238–247.

Cordeiro ZJM and Matos AP (2003) Impact of *Mycosphaerella spp*. in Brazil. In: Jacome et al. (eds.). Mycosphaerella leaf spot diseases of bananas: present status and outlook. proceedings of the workshop on Mycosphaerella leaf spot diseases. Montpellier, France, p. 91–97.

Cordeiro ZJM, Matos AP, Kimati H (2005) Doenças da bananeira. In: Kimati et al. (eds). Manual de fitopatologia: Doenças das Plantas Cultivadas. São Paulo, Brazil, p. 99–117.

Cordeiro et al (2004) Doenças e Métodos de Controle. In: Borges, A.L., Souza, L.S. (eds) O Cultivo da Bananeira. Cruz das Almas, Bahia, Brasil, p. 146–182.

Cordeiro et al. (2011) Metodologias para Manuseio de *Mycosphaerella musicola* em Laboratório. Embrapa mandioca e fruticultura. Cruz das Almas-BA, Brazil.

Daub et al. (2005) Photoactivated perylenequinone toxins in fungal pathogenesis of plants. FEMS Microbiology Letters 252:197–206.

Dold et al. (2008) Musa in Shaded Perennial Crops - Response to Light Interception. Conference on international research on food security, natural resource management and rural development. Stuttgardt, Germany, p.1–4.

Emebiri LC and Obiefuna JC (1992) Effects of leaf removal and intercropping on the incidence and severity of black Sigatoka disease at the establishment phase of plantains (*Musa spp.* AAB). Agriculture, ecosystems and environment 39:213–219.

Etebu et al. (2005) Effect of light and sealing pattern on sporulation and growth of *Mycosphaerella fijiensis*. In: Vézina, A. The international journal on banana and plantain, 14:24–25.

Food and Agriculture Organization of the United Nations Agricultural Data base - FAO. 2011. Avaliable at: http://www.fao.org. Accessed 16 November 2011.

Favreto et al. (2007) Sigatoka Negra, fatores de ambiente e sistemas agroflorestais em bananais do Rio Grande do Sul, Brazil. Pesquisa Agropecuária Gaúcha, Porto Alegre 13:95–104.

Hanada et al. (2002) Esporulação de *Mycosphaerella fijjiensis* em diferentes meios de cultura. Fitopatologia Brasileira 27:170–173.

Jácome LH (2002) Population biology and epidemiology. In: Jacome et al. (eds.). Mycosphaerella leaf spot diseases of bananas: present status and outlook. Proceedings of the workshop on mycosphaerella leaf spot diseases. Montpellier, France, p.107-110.

Jácome et al. (1991) Effect of Temperature and Relative Humidity on Germination and Germ tube Development of *Mycosphaerella fijiensis var difformis*. Phytopathology 81:1480–1485.

Jones DR (2002) The distribution and importance of the *Mycosphaerella* leaf spot diseases of banana. In: Jacome et al. (eds.). Mycosphaerella leaf spot diseases of bananas: present status and outlook. Proceedings of the workshop on Mycosphaerella leaf spot diseases. Montpellier, France, p. 25–41.

Lepoivre et al. (2002) Banana – *Mycosphaerella fijiensis* interactions. In: Jacome et al. (eds.). Mycosphaerella leaf spot diseases of bananas: present status and outlook. Proceedings of the workshop on Mycosphaerella leaf spot diseases. Montpellier, France, p. 151–159.

Montarroyos et al. (2007) Efeitos de meio de cultura, fontes de carbono e nitrogênio, pH e regime luminoso no crescimento de *Mycosphaerella musicola*. Summa Phytopathologica 33:86–89.

Norgrove L and Hauser S (2013) Black leaf streak disease and plantain fruit characteristics as affected by tree density and biomass management in a tropical agroforestry system. Agroforestry Systems 87:349. doi:10.1007/s10457-012-9555-z.

Norgrove L (1998) Musa in multistrata systems: focus on shade. Infomusa 7:17–22.

Norgrove et al. (2012) Tackling black leaf streak disease and soil fertility constraints to enable the expansion of plantain production to grassland in the humid tropics. International Journal of Pest Management 58:175–181. doi: 10.1080/09670874.2012.676218.

Rocha et al. (2012) Temporal Progress of Yellow Sigatoka and Aerobiology of *Mycosphaerella musicola* Spores. J Phytopathol 160:277–285.

Rosa RCT and Menezes M (2001) Caracterização patogênica, fisiológica e morfológica de *Pseudocercospora musae*. Fitopatologia Brasileira 26:141–147.

Sepúlveda et al. (2009) The presence and spectrum of light influences the in vitro conidia production of *Mycosphaerella fijiensis* causal agent of black Sigatoka. Australasian Plant Pathology 38:514–517.

Stierle et al. (1991) The phytotoxins of *Mycosphaerella fijiensis*, the causative agent of Black Sigatoka disease of bananas and plantains. Experientia 47:853. doi:10.1007/BF01922472.

Vivan JL (2002) Bananicultura em Sistemas Agroflorestais no Litoral Norte do RS. Agroecologia e Desenvolvimento Rural Sustentável 3:17–26.

